# Tiling light sheet selective plane illumination microscopy using discontinuous light sheets

**DOI:** 10.1101/378232

**Authors:** Liang Gao

## Abstract

Tiling light sheet selective plane illumination microscopy (TLS-SPIM) improves 3D imaging ability of SPIM by using a real-time optimized tiling light sheet. However, the imaging speed decreases, and size of the raw image data increases proportionally to the number of tiling positions in TLS-SPIM. The decreased imaging speed and the increased raw data size could cause significant problems when TLS-SPIM is used to image large specimens at high spatial resolution. Here, we present a novel method to solve the problem. Discontinuous light sheets created by scanning coaxial beam arrays synchronized with camera exposures are used for 3D imaging to decrease the number of tiling positions required at each image plane without sacrificing the spatial resolution. We investigate the performance of the method via numerical simulation and discuss the technical details of the method.

## Introduction

The 3D imaging ability of selective plane illumination microscopy (SPIM), i.e. light sheet microscopy, relies on the intensity profile of the light sheet used for 3D imaging. The thickness, light confinement ability, and size of the light sheet determine the axial resolution, optical sectioning ability, and field of view (FOV) of SPIM respectively. [1,2] In order to improve the 3D imaging ability of SPIM on large specimens, tremendous efforts have been spent to optimize the light sheet intensity profile in the above aspects with the objective of confining the excitation light near the detection focal plane over a long distance as much as possible.[3–7]

Unfortunately, a fundamental trade off exists among the above properties of a light sheet due to the diffraction of light. The excitation light is always less confined as the light sheet size increases, which makes it extremely difficult to image large specimens at high spatial resolution and signal to noise ratio (SNR) using SPIM. Methods other than optimizing the light sheet intensity profile were developed to solve the problem [8–13]. Among these methods, an effective approach is to quickly move the light sheet axially within the image plane during imaging, so that high spatial resolution and good optical sectioning ability can be maintained in a much larger FOV compared to the size of the light sheet itself [14–19]. Tiling light sheet selective plane illumination microscopy (TLS-SPIM) is a method using this strategy to improve the 3D imaging ability of SPIM on large specimens. [18,19]

In TLS-SPIM, a large field of view (FOV) is imaged by tiling a short but thin light sheet at multiple positions within the detection focal plane and taking a camera exposure at each light sheet tiling position. It has been demonstrated that TLS-SPIM is capable to image large multicellular specimens of different sizes, ranging from live embryonic specimens to cleared tissue specimens, at high spatial resolution [19,20]. In addition, TLS-SPIM allows the optimization of the tiling light sheet intensity profile and the tiling position in less than a millisecond, so that the 3D imaging ability of the microscope can be optimized in real-time based on the biological specimen and process being imaged.

Despite the improved 3D imaging ability of TLS-SPIM, the additional camera exposures required in the light sheet tiling process cause major problems. The imaging speed decreases, and the size of the raw image data increases proportionally to the number of tiling positions, i.e. the number of camera exposures required per image plane. Although these problems are less of an issue when the number of tiling positions or sample size is small, it could be truly troubling when a large specimen needs to be imaged by TLS-SPIM at a high spatial resolution, which requires a large number of tiling positions per image plane. For example, when TLS-SPIM is used to image a cleared tissue specimen of dozens of cubic millimeters at micron level spatial resolution, the total imaging time could be extended by a few hours or more compared to a conventional non-tiling condition, despite the improved spatial resolution. Meanwhile, hundreds of gigabytes, even dozens of terabytes, additional raw image data, are created by the additional camera exposures, and must be collected, which produce a heavy burden to the often limited data collection bandwidth and the data storage space.

In this research, we developed a novel method to address the above problems of TLS-SPIM. Discontinuous tiling light sheets, created by scanning coaxial beam arrays that are synchronized with the rolling shutter of the detection sCMOS camera, are used in TLS-SPIM to image a large FOV, by which the imaging speed of TLS-SPIM can be increased, and the raw image data size can be deceased proportionally to the number of the coaxial beams contained in the coaxial beam array. Here, we investigate the method and evaluate its performance via numerical simulation.

## Methods

In order to increase the imaging speed of TLS-SPIM and produce less raw image data, the number of tiling positions and camera exposures must be decreased to image the same FOV. In other words, a larger effective area must be imaged at each light sheet tiling position with each camera exposure. Thus, the tiling light sheet must be enlarged along the light propagation direction. Meanwhile, the thickness and light confinement ability of the light sheet must remain the same to ensure that the same spatial resolution and optical sectioning ability can be maintained.

Using a “non-diffracting” light sheet, such as the Bessel light sheet or Lattice light sheet, is a possible solution to address the problem. However, the excitation light is still less confined by a “non-diffracting” light sheet when its size increases, which reduces the optical sectioning ability of SPIM significantly. Therefore, we decided to seek a different approach to solve the problem. Instead of using a conventional light sheet with a continuous intensity profile, multiple light sheets that are separated far enough in the light propagation direction are used and tiled simultaneously for 3D imaging in TLS-SPIM. It can also be considered as using a discontinuous light sheet with multiple waists instead of a single continuous waist (Fig. 1), as the light sheet doesn’t need to be continuous in TLS-SPIM. There are at least two ways to use discontinuous light sheets in TLS-SPIM for 3D imaging. First, the entire FOV can be imaged by tiling the same discontinuous light sheet at different positions in the FOV, which is the same as that in regular TLS-SPIM (Fig. 2a). Second, the entire FOV can be imaged by using multiple different discontinuous light sheets with different waist numbers and positions to compensate each other, by which the entire FOV can also be imaged after a few cycles (Fig. 2b).

**Figure 1.**
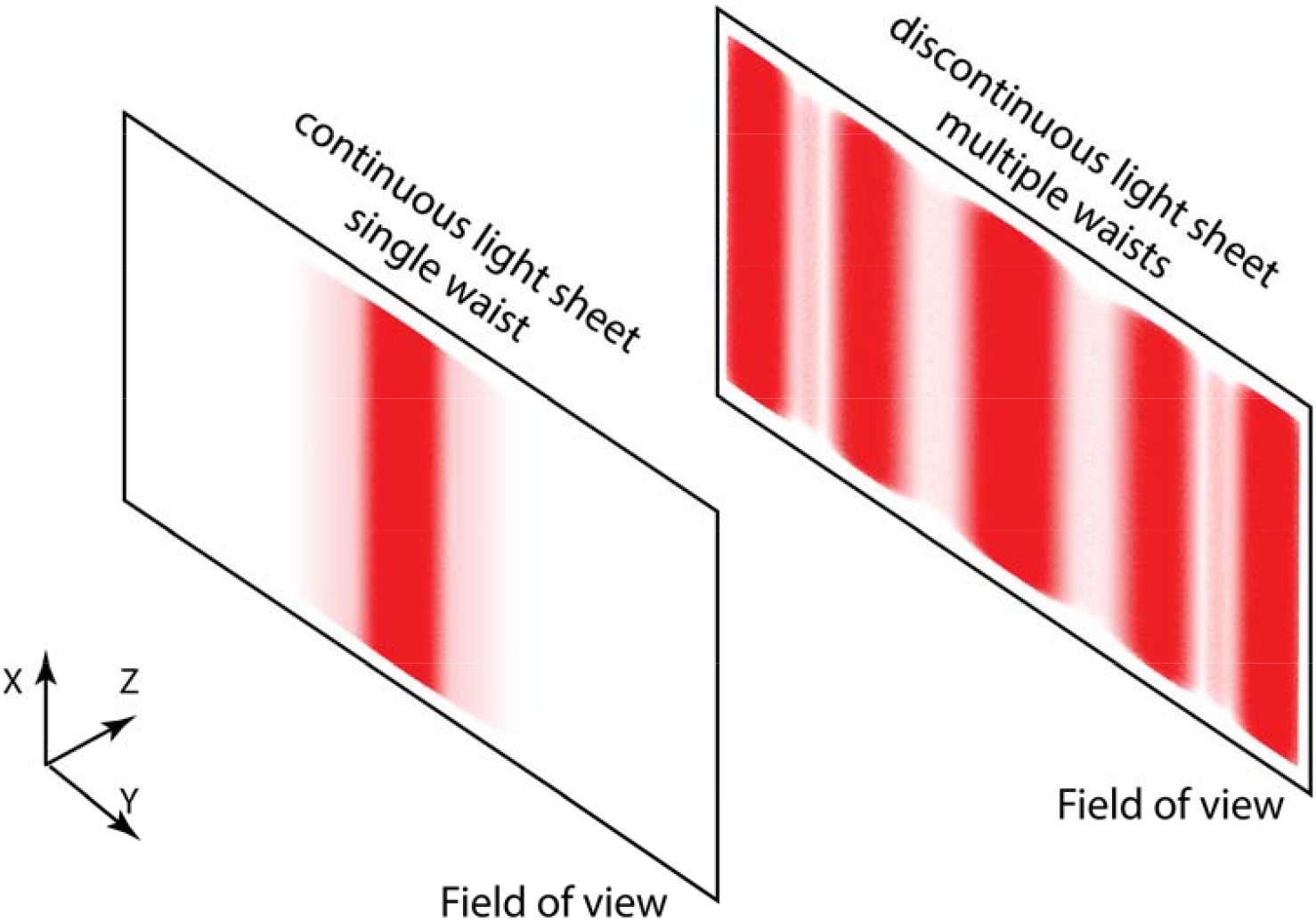
The concept of the method. A discontinuous light sheet with multiple waists allows a larger area being imaged at each tiling position compared to a continuous light sheet with a single waist, which improves the imaging speed and decreases the raw image data size of TLS-SPIM at the same time.

**Figure 2.**
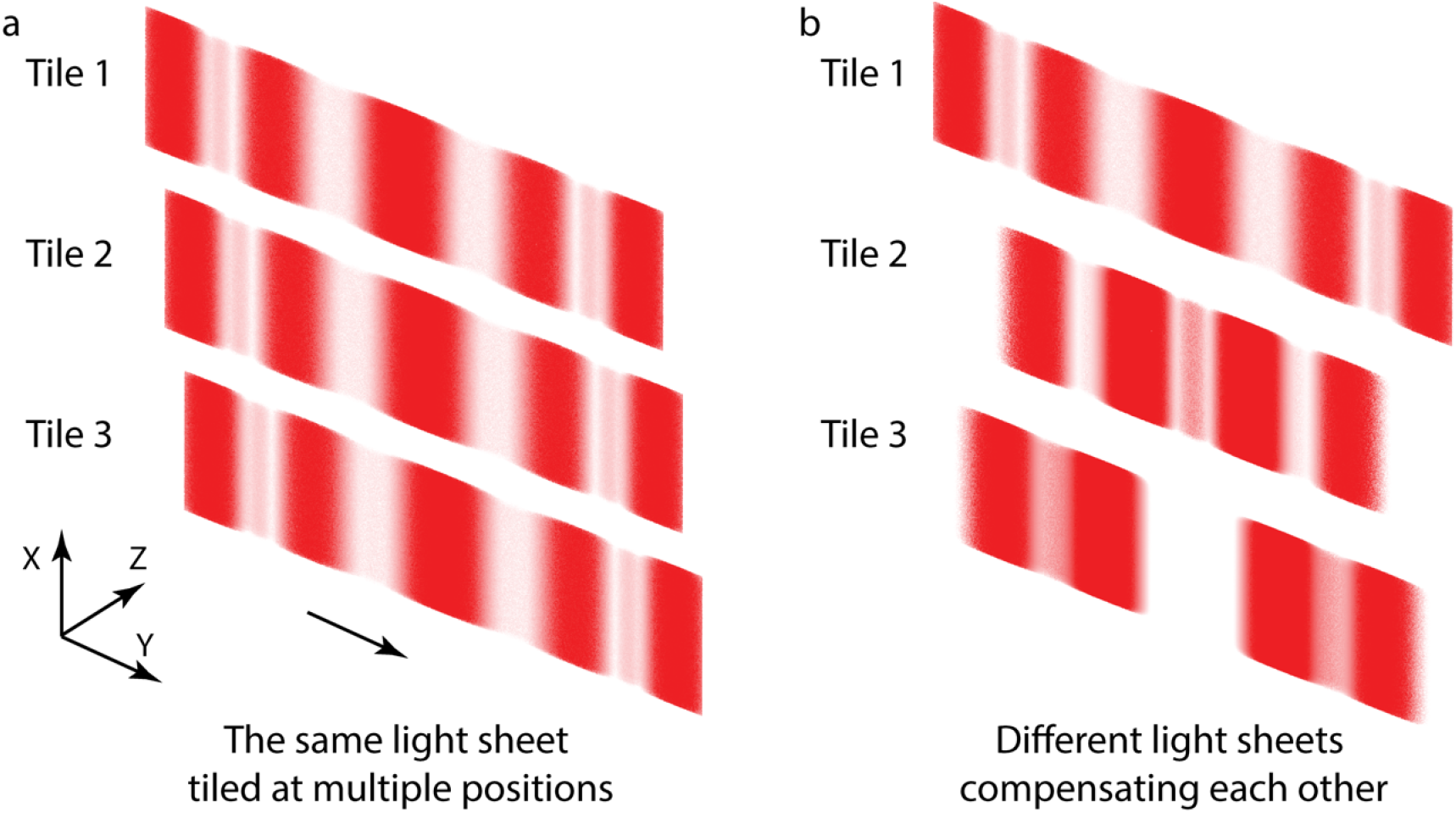
Two methods to implement discontinuous light sheets in TLS-SPIM for 3D imaging, (a) A field of view can be imaged by tiling the same discontinuous light sheet at multiple positions. (b) A field of view can be imaged by using multiple different discontinuous light sheets that compensate each other.

See the light, we investigated the optical designs to conduct the above operations in TLS-SPIM. In the TLS-SPIM microscope reported previously [19], either one or two binary SLMs are used to generate and tile light sheets of different dimensions for 3D imaging. Although the light sheet tiling process can also be realized by other optical devices, such as a focal variable lens, digital mirror device (DMD), deformable mirror, continuous phase SLM or a mechanical scanning device, the binary SLM has signification advantages compared to others due to its fast response speed, accurate phase modulation ability, control flexibility, robustness and the lack of mechanical movements, so does the image registration. The only major disadvantage of the binary SLM is perhaps a relative low laser power efficiency caused by the undesired diffraction orders generated by the device, which can be easily solved by using a high power laser source. Thus, we decided to proceed with optical designs of using binary SLMs to implement discontinuous light sheets in TLS-SPIM.

A discontinuous light sheet can be obtained by scanning a coaxial beam array, and the effective size of such a discontinuous light sheet is proportional to the number of coaxial beams contained in the beam array. A coaxial beam array can be created easily by adding a DOE element into the previous TLS-SPIM system at a plane conjugated to the rear pupil of the excitation objective, and the binary SLM can be used as a focal variable device to tile the coaxial beam array, so as to the discontinuous light sheet. However, a DOE element can only generate a coaxial beam array with a fixed beam number and period, which reduces the flexibility of TLS-SPIM significantly. In addition, a DOE element usually works differently for different excitation wavelengths, which makes it difficult to maintain the same imaging quality for different excitation wavelengths. Therefore, we seek solutions to generate coaxial beam arrays using the binary SLM used in TLS-SPIM, so that the microscope can generate coaxial beam arrays of different beam numbers and periods for different excitation wavelengths, tile the beam array, and switch between different beam arrays quickly to optimize the imaging performance.

Methods of generating coaxial beam arrays using binary optical devices has been studied extensively. A method that is modified from the designing of Dammann gratings was developed in our research to generate binary phase maps for the binary SLM to create coaxial beam arrays of different dimensions [21]. As shown in Fig. 3, coaxial beam arrays consisted of 1 to 4 Gaussian beams (0.07 excitation NA) with different beam array periods were generated by applying different binary phase maps to a binary SLM. Nevertheless, it must be aware that the choices of the beam array periods and beam numbers are still limited due to the limited resolution of the binary SLM, despite it is much more flexible than a fixed DOE element.

**Figure 3.**
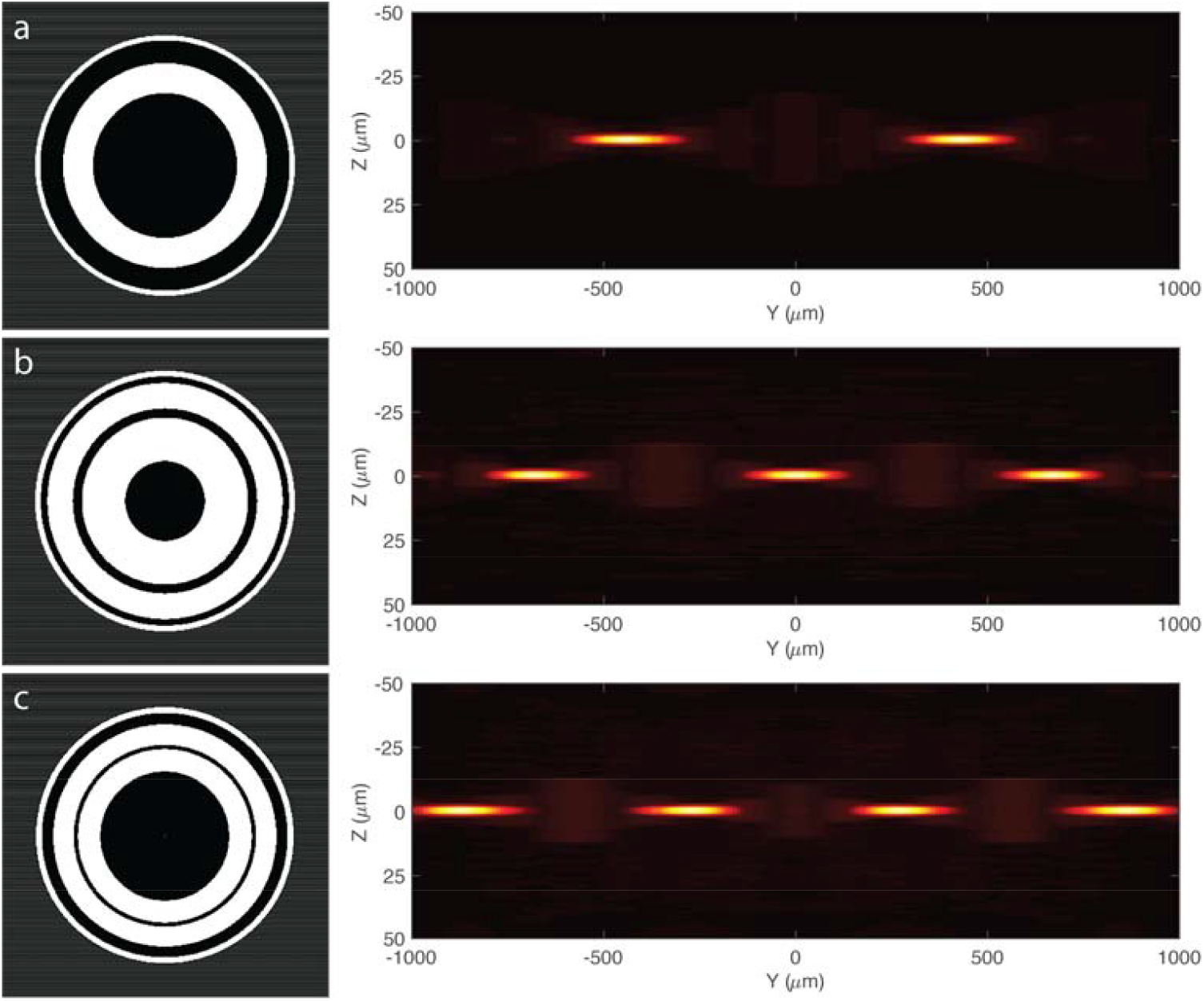
Binary phase maps and the corresponding coaxial beam arrays that contain different number of beams and beam array periods. Excitation NA_od_=0.08, NA_id_=0.02.

Next, the coherent beam array needs to be tiled along the light propagation direction to image the entire FOV. As discussed above, there are two ways to perform the light sheet tiling process. One is to tile the same coaxial beam array at different positions in the same way as that in regular TLS-SPIM. It can be achieved by superimposing a tilted spherical phase to the the circular Dammann grating used to generate the coaxial beam array and resetting the phase values to 0 and *π* (Fig. 4). A similar tiling process can also be realized by moving the sample if the specimen is fixed and the imaging speed is not a concern. Otherwise, moving the sample is generally less preferred in most applications.

**Figure 4.**
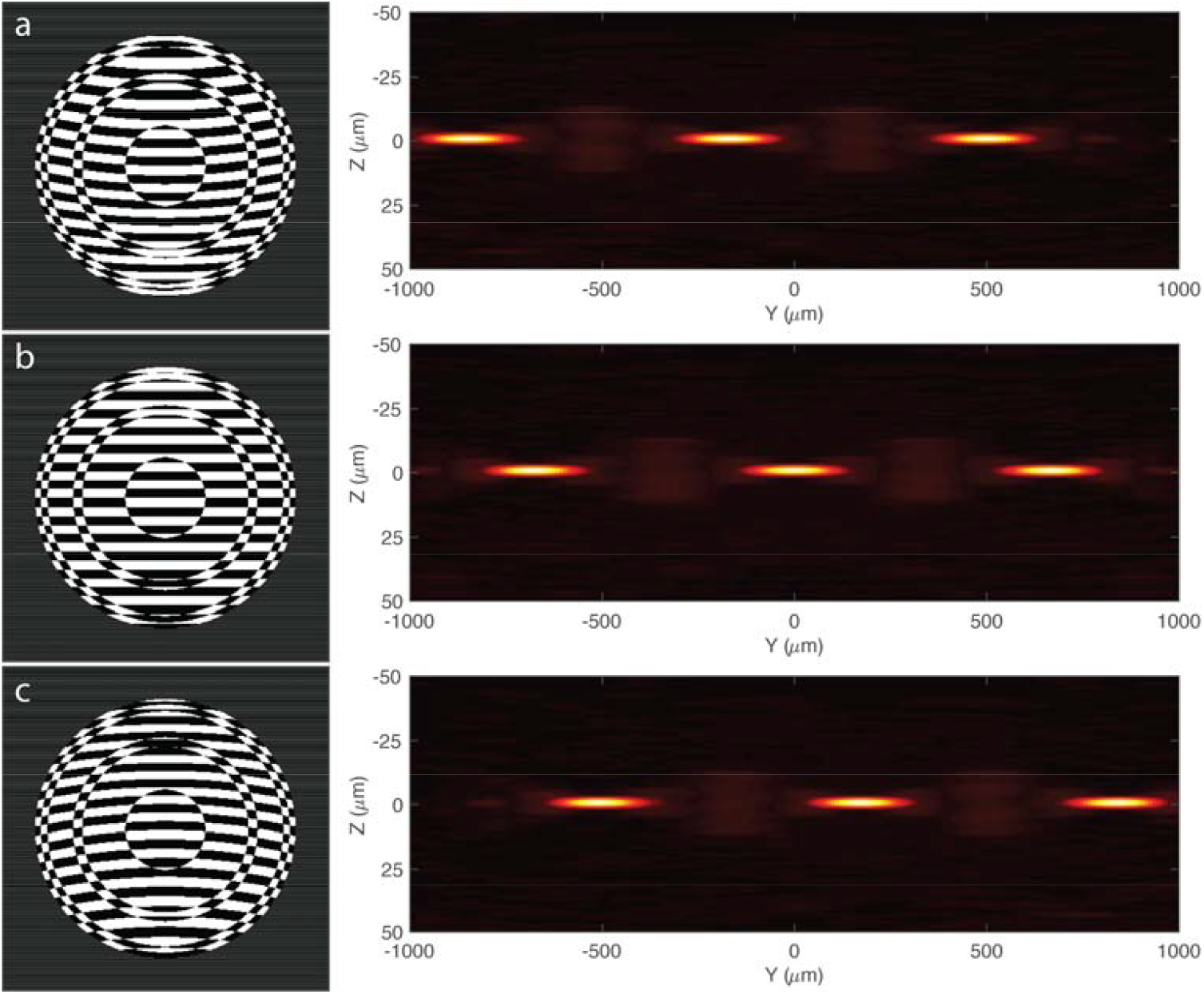
Binary phase maps used to create and tile the same three beam coaxial beam array at different positions within the field of view. Excitation NA_od_=0.08, NA_id_=0.02.

Instead of tiling the same light sheet, the other way to image the entire FOV in TLS-SPIM is to use multiple discontinuous light sheets with different waist positions, that compensate each other. For example, a FOV can be imaged by using three discontinuous light sheets, which are created by scanning a five beam coaxial beam array, and two four beam arrays respectively (Fig. 5). An additional advantage of the method is that more diffraction orders created by the binary SLM are used for imaging, so that the laser efficiency is improved significantly compared to the previous method.

**Figure 5.**
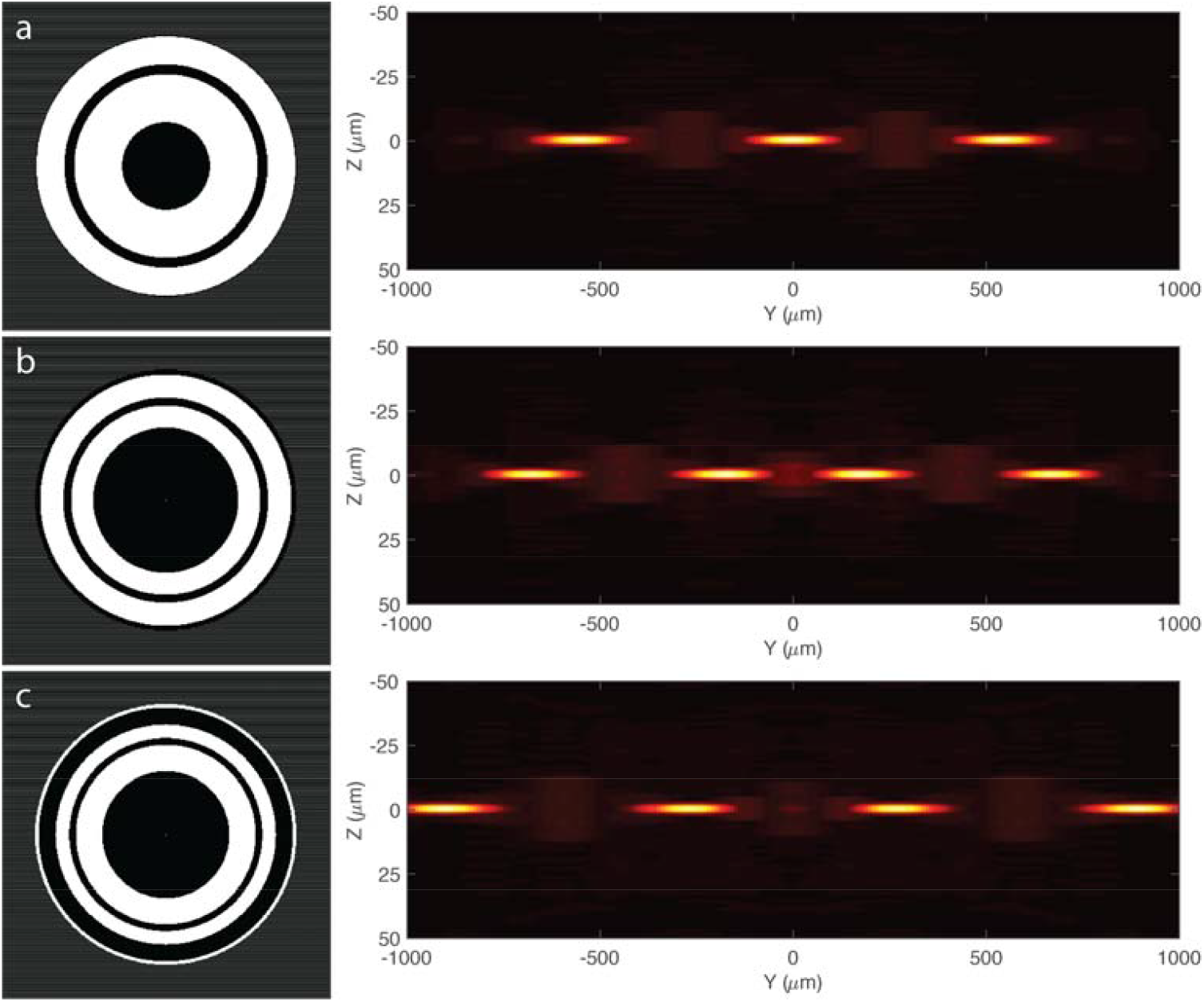
Binary phase maps used to generate three different coaxial beam arrays with beam positions compensate each other. Excitation NA_od_=0.08, NA_id_=0.02.

Nevertheless, the excitation coaxial beam array needs to be scanned to create a virtual discontinuous light sheet at each tiling position, while the crosstalk between different beams of a coaxial beam array causes a problem when the beam array is scanned to created a discontinuous light sheet. The off-focus light of the beam array adds up together during the beam scanning process and introduces additional off-focus excitation at the waist positions of the discontinuous light sheet, which the same as what happens with a “non-diffracting” light sheet. Obviously, the additional off-focus excitation reduces the optical sectioning capability of the microscope. As shown (Fig. 6), the off-focus excitation energy increases as the beam array period decreases and the beam number increases. Indeed, the diffraction of light always dominates the tradeoff between the thickness, light confinement ability and size of a light sheet. The light confinement ability of a light sheet decreases as the usable portion of the light sheet increases, regardless the exact intensity profile. Despite the similarity between a discontinuous light sheet and a “non-diffracting” light sheet, that both of them extend the usable portion of a light sheet by sacrificing the excitation light confinement ability, there is a significant difference between them, which is the distribution of the unconfined excitation light outside the detection focal plane. Figure 6 and Figure 7 show the intensity profile differences between discontinuous light sheets and Bessel light sheets that are capable to image the same FOV at the same theoretical axial resolution. As shown, the off focus excitation light is spread much further from the detection focal plane in discontinuous continuous light sheets created by scanning a coaxial beam array compared to the Bessel light sheets.

**Figure 6.**
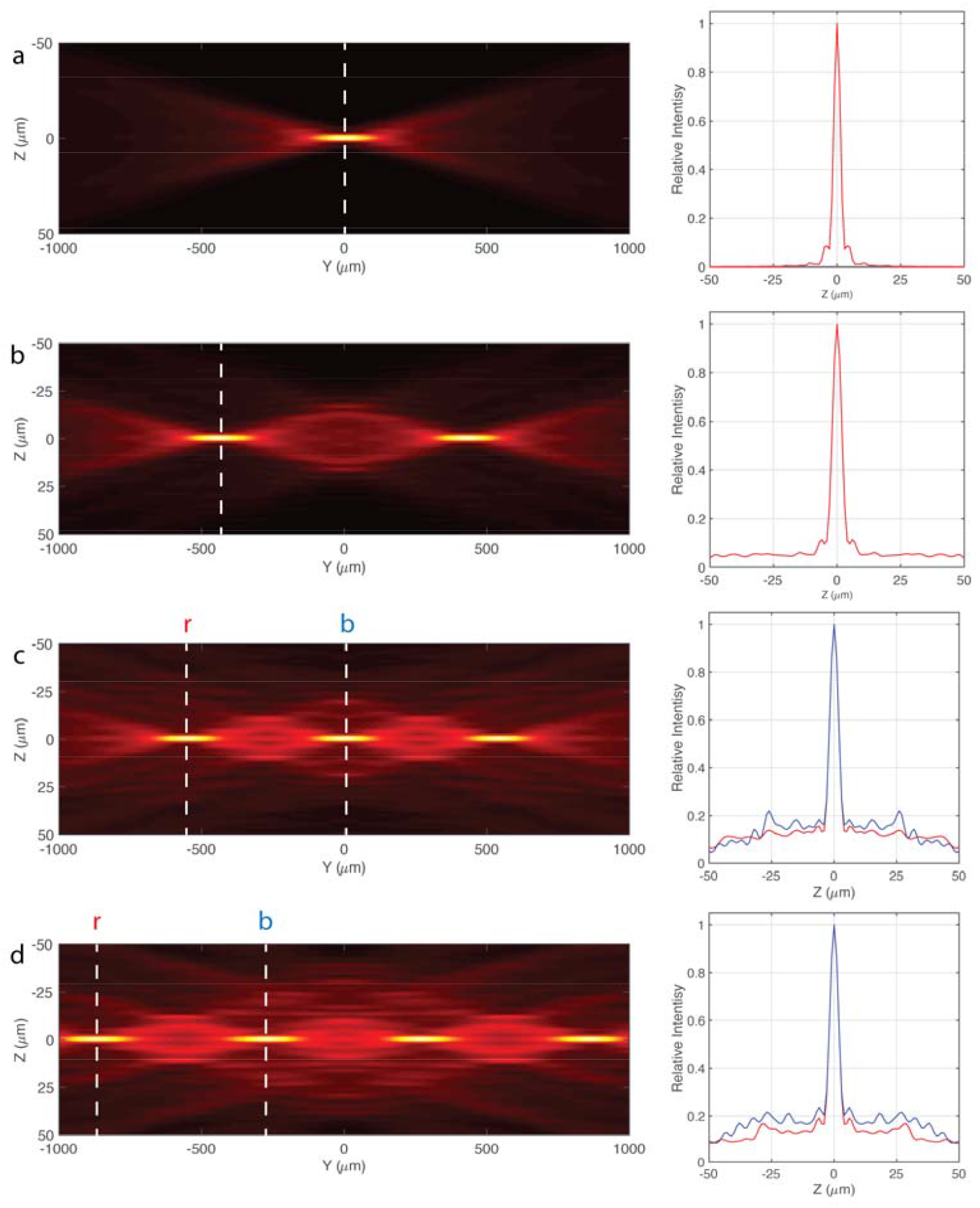
YZ max projection and the intensity profile of discontinuous light sheets created by scanning coaxial beam arrays that contain two to four coaxial beams. Excitation NA_od_=0.08, NA_id_=0.02.

**Figure 7.**
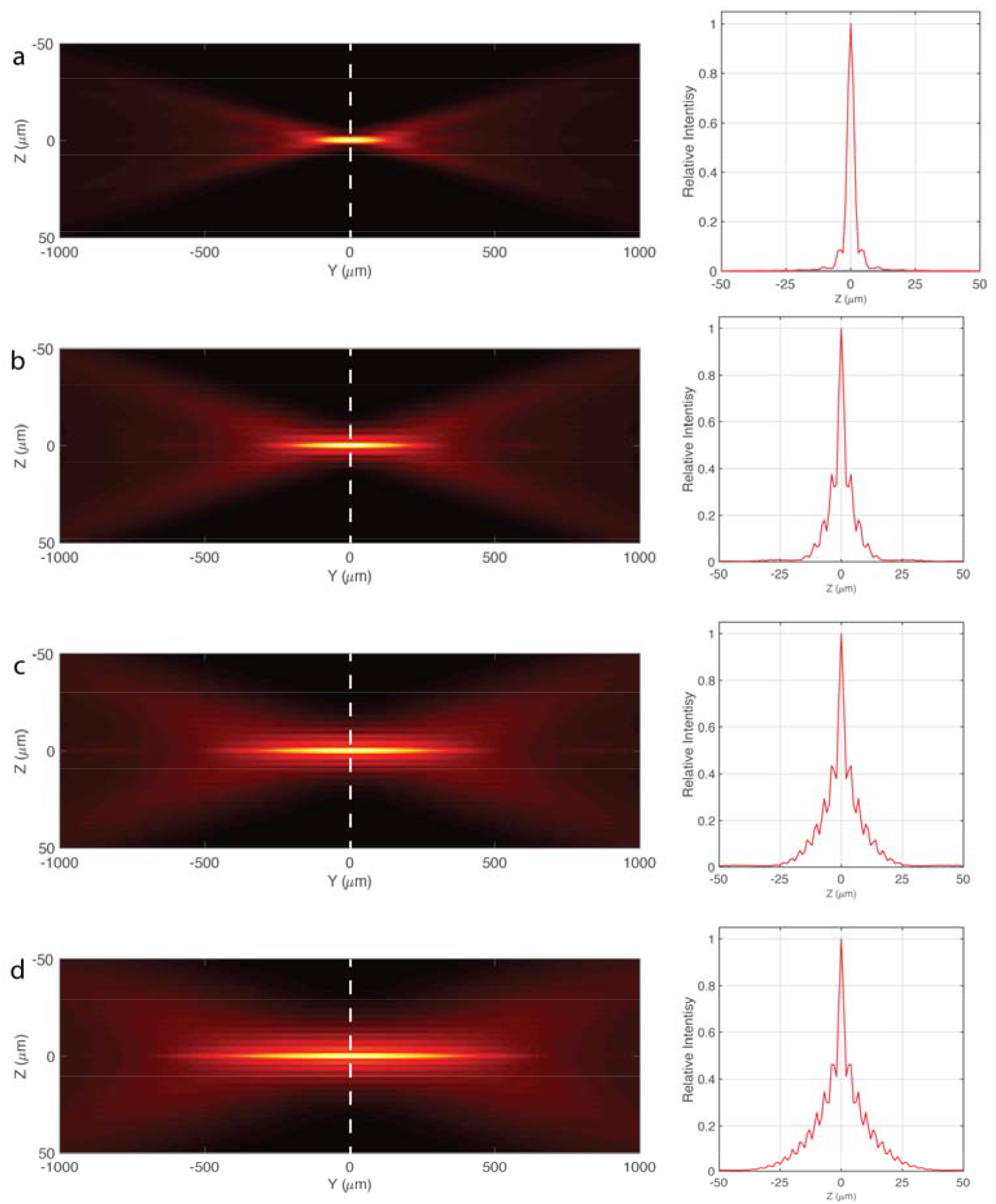
YZ max projection and the intensity profile of Bessel light sheets that offer comparable field of view and axial resolution to the discontinuous light sheet in Fig. 6. Excitation NA=0.08, NA_id_=0.02 in a, NA_od_=0.08, NA_id_=0.057 in b, NA_od_=0.08, NA_id_=0.065 in c and NA_od_=0.08, NA_id_=0.069 in d.

In light sheet microscopy, the off-focus fluorescence background produced by a virtual light sheet created by a scanning beam can be suppressed by using a sCMOS camera for fluorescence detection and operating the sCMOS camera in the light sheet readout mode [22,23]. In the light sheet readout mode of a sCMOS camera, the exposure and readout of each pixel row is synchronized with the scanning coaxial beam array, which results in a slit confocal detection effect rather than the wide field detection in normal light sheet microscopy, so that the majority of the fluorescence background created by the off-focus excitation light is rejected by the camera. Obviously, the off-focus fluorescence background created by discontinuous light sheets can be rejected much easier because the confocal slit detection is more effective in rejecting the fluorescence background that is further away from the detection focal plane.

Next, we studied the effectiveness of the confocal slit detection in rejecting the off-focus fluorescence background created by discontinuous light sheets with different detection numerical apertures (NA) and the confocal slit widths. Fig. 8–9 show the simulated equivalent light sheet intensity profile of the discontinuous light sheets in Fig. 6 with 3.5-13.5 μm confocal slit detection at 0.25, 0.37 and 0.5 detection. We also simulated the equivalent light sheet intensity profiles of Bessel light sheets in Fig. 7 that offer comparable theoretical spatial resolution and FOV. Clearly, the confocal slit detection is an effective method to suppress the off-focus fluorescence background. It behaves very similar with different detection NA, but thinner confocal slits reject the off-focus background much more effectively. Meanwhile, it is obvious that the further the off-focus background is away from the detection focal plane, the more it is suppressed by the confocal slit detection, which is a key advantage of the discontinuous light sheet over “non-diffracting” light sheets, such as the Bessel light sheet. As a result, a large area can be imaged at each tiling position in TLS-SPIM using discontinuous light sheets together with confocal slit detection, by which the imaging speed of TLS-SPIM can be increased and the size of raw imaged data can be decreased significantly with little loss on the optical sectioning capability.

**Figure 8.**
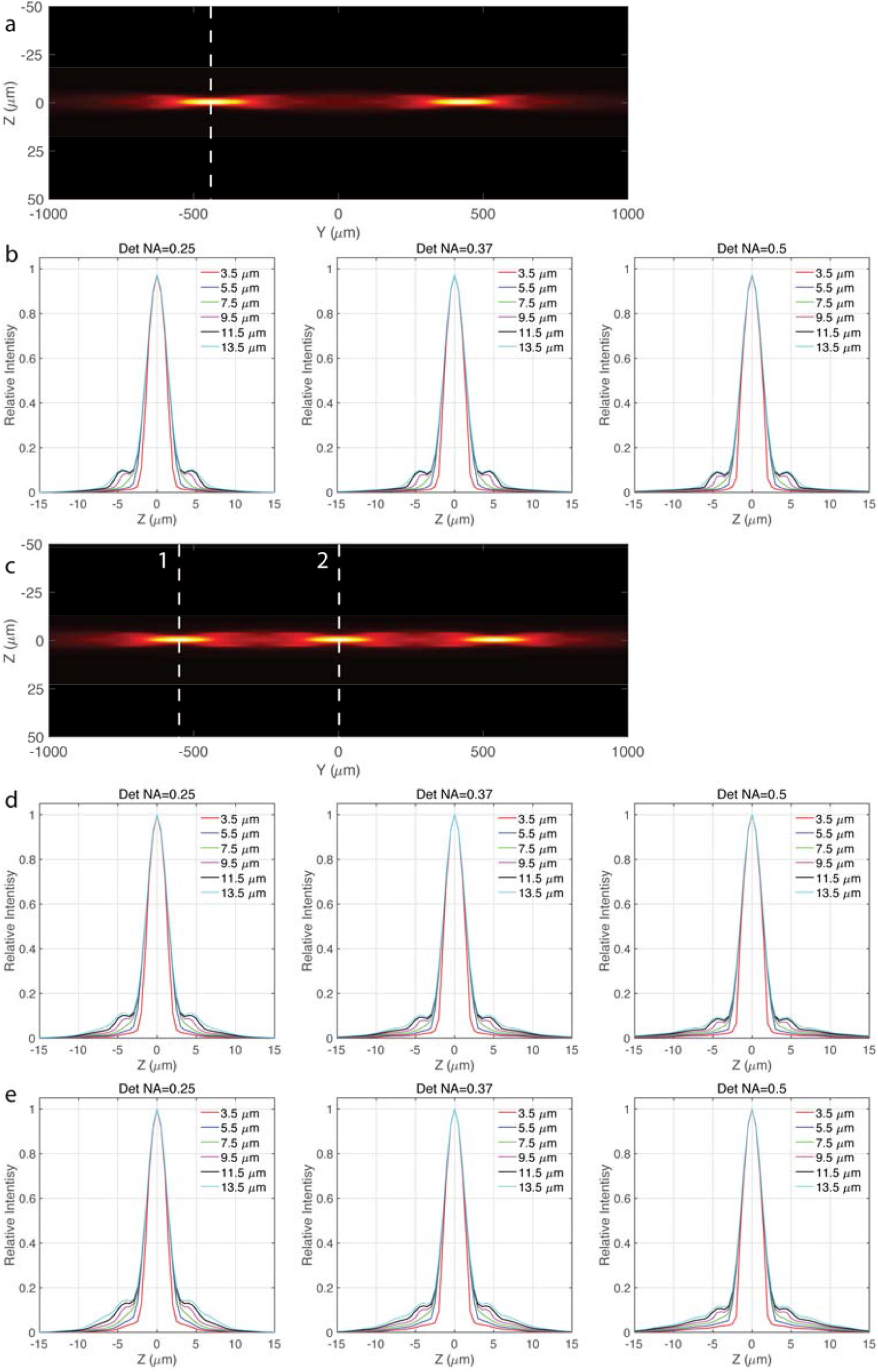
Equivalent light sheets of the discontinuous light sheets 6b and 6c with confocal slit detection, (a) The YZ projection of the equivalent light sheet of the discontinuous light sheet 6b with 7.5 μm confocal slit and 0.37 detection NA. (b) The intensity profile of the equivalent light sheet of the discontinuous light sheet 6b with different confocal slit width and detection NA. (c) The YZ projection of the equivalent light sheet of the discontinuous light sheet 6c with 7.5 μm confocal slit and 0.37 detection NA. (d) The intensity profile of the equivalent light sheet of the discontinuous light sheet 6c at position 1 with different confocal slit width and detection NA. (e) The intensity profile of the equivalent light sheet of the discontinuous light sheet 6c at position 2 with different confocal slit width and detection NA.

**Figure 9.**
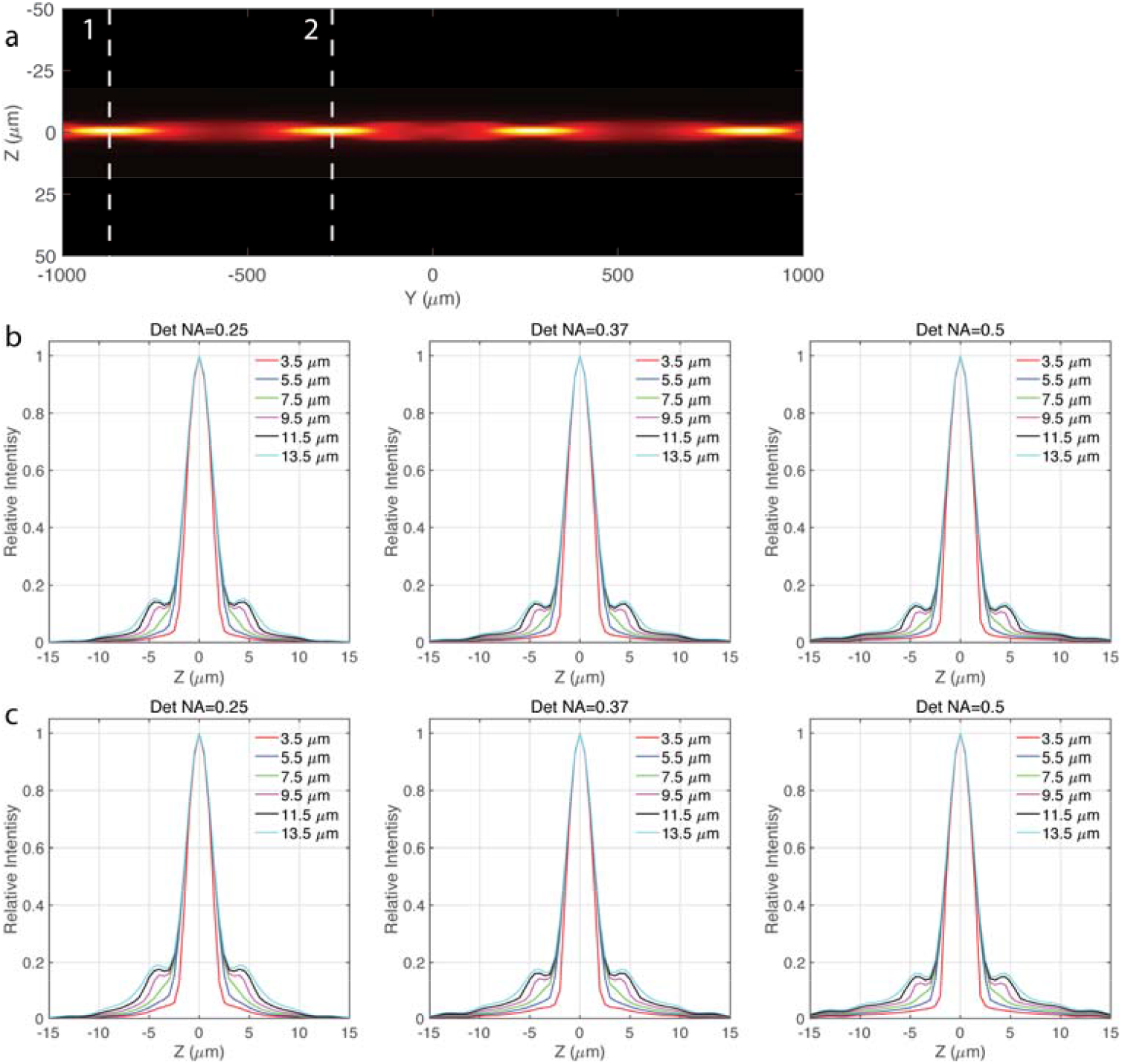
(a) The YZ projection of the equivalent light sheet of the discontinuous light sheet 6d with 7.5 μm confocal slit and 0.37 detection NA. (b) The intensity profile of the equivalent light sheet of the discontinuous light sheet 6d at position 1 with different confocal slit width and detection NA. (c) The intensity profile of the equivalent light sheet of the discontinuous light sheet 6d at position 2 with different confocal slit width and detection NA.

**Figure 10.**
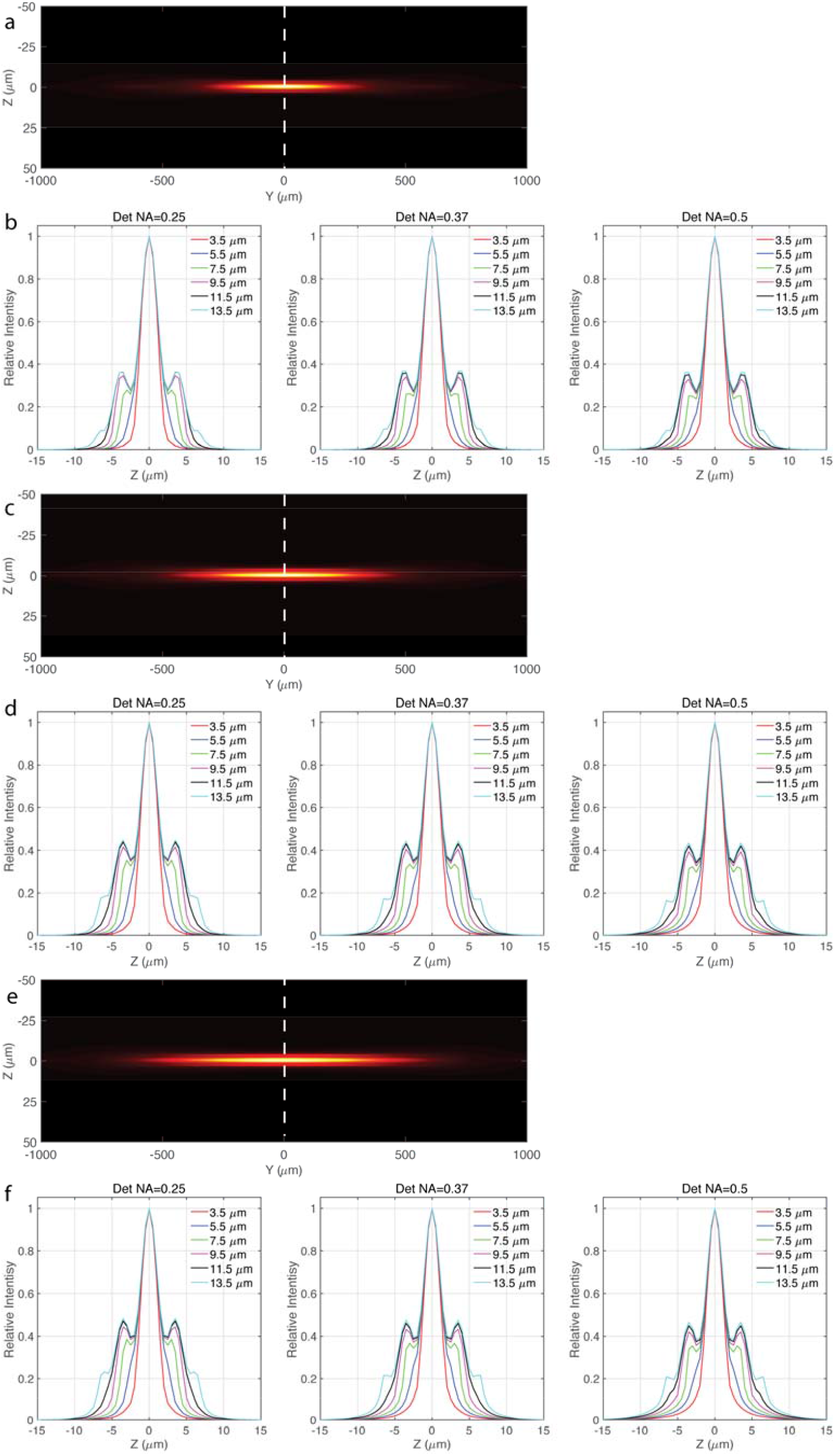
Equivalent light sheets of the Bessel light sheets 7b, 7c and 7d with confocal slit detection, (a) The YZ projection of the equivalent light sheet of the Bessel light sheet 7b with 7.5 μm confocal slit and 0.37 detection NA. (b) The intensity profile of the equivalent light sheet of the Bessel light sheet 7b with different confocal slit width and detection NA. (c) The YZ projection of the equivalent light sheet of the Bessel light sheet 7c with 7.5 μm confocal slit and 0.37 detection NA. (d) The intensity profile of the equivalent light sheet of the Bessel light sheet 7c with different confocal slit width and detection NA. (e) The YZ projection of the equivalent light sheet of the Bessel light sheet 7d with 7.5 μm confocal slit and 0.37 detection NA. (f) The intensity profile of the equivalent light sheet of the Bessel light sheet 7d with different confocal slit width and detection NA.

## Conclusions

In summary, we present a novel method to increase the imaging speed and decrease the raw data size of TLS-SPIM. Discontinuous lights sheets created by scanning coaxial beam arrays synchronized with the camera exposure are used for 3D imaging. We study the method and evaluated its performance via numerical simulations. We show that the implementation of discontinuous light sheets together with the confocal slit detection mode in TLS-SPIM allow the same FOV being imaged with less light sheet tiling positions, i.e. camera exposures, while maintain the same spatial resolution and optical sectioning ability. The new method can improve the imaging speed and reduce the image data size by several times up to the discontinuous light sheet be used, which could bring significant benefits when TLS-SPIM is used to image large specimens at high spatial resolution.

